# TRACE: transcription factor footprinting using chromatin accessibility data and DNA sequence

**DOI:** 10.1101/801001

**Authors:** Ningxin Ouyang, Alan P. Boyle

## Abstract

Transcription is tightly regulated by cis-regulatory DNA elements where transcription factors can bind. Thus, identification of transcription factor binding sites is key to understanding gene expression and whole regulatory networks within a cell. The standard approaches for transcription factor binding sites (TFBSs) prediction such as position weight matrices (PWMs) and chromatin immunoprecipitation followed by sequencing (ChIP-seq) are widely used but have their drawbacks such as high false positive rates and limited antibody availability, respectively. Several computational footprinting algorithms have been developed to detect TFBSs by investigating chromatin accessibility patterns, but also have their limitations. To improve on these methods, we have developed a footprinting method to predict Transcription factor footpRints in Active Chromatin Elements (TRACE). Trace incorporates DNase-seq data and PWMs within a multivariate Hidden Markov Model (HMM) to detect footprint-like regions with matching motifs. Trace is an unsupervised method that accurately annotates binding sites for specific TFs automatically with no requirement on pre-generated candidate binding sites or ChIP-seq training data. Compared to published footprinting algorithms, TRACE has the best overall performance with the distinct advantage of targeting multiple motifs in a single model.

## Introduction

Identification of the cis-regulatory elements where transcription factors (TFs) bind remains a key goal in the deciphering of transcriptional regulatory circuits. Standard approaches to identify the set of active transcription factor binding sites (TFBSs) include the use of position weight matrices (PWMs) (Stormo et al. 1982) and ChIP-seq (Barski et al. 2007). While these methods have had success, they individually suffer from drawbacks that limits their usefulness. PWMs, while able to identify high-resolution binding sites, are prone to extremely high false positive rates in the genome. On the other hand, while ChIP-seq measurements of binding is highly specific and have a significantly reduced false positive rate, their resolution is comparatively low, they are labor intensive, and they depend on suitable antibodies which are only available for a limited number of TFs. Newer experimental techniques for identification DNA-bound proteins binding sites such as ChIP-exo (Rhee and Pugh 2012) and CUT&RUN (Skene and Henikoff 2017) have the advantage of high resolution and cost efficiency, but still share the same disadvantages as ChIP-seq including labor demanding and limited antibody availability.

To complement these approaches, another experimental method has been developed using data from high-throughput sequencing after DNase I digestion (DNase-seq) (Boyle et al. 2008). These DNase-seq data identify stretches of open regions of chromatin where DNase I cuts at a higher frequency. Within these regions, TFBSs can be identified at nucleotide resolution by searching for footprint-like regions with low numbers of DNase I cuts embedded in peaks with high numbers of cuts.

Hesselberth et al. (2009) first proposed a DNase-seq signal based computational method to detect footprints at base pair resolution in *Saccharomyces cerevisiae*. Since then, several computational footprinting algorithms have been developed to detect TFBSs by investigating chromatin accessibility patterns which can be categorized as *de novo* (the Boyle method, DNase2TF, HINT, PIQ and Wellington) and motif-centric (DeFCoM, BinDNase, CENTIPEDE, FLR) (Boyle et al. 2011; Gusmao et al. 2016; Kähärä and Lähdesmäki 2015; Piper et al. 2013; Pique-Regi et al. 2011; Quach and Furey 2016; Sherwood et al. 2014; Sung et al. 2014; Yardımcı et al. 2014). *De novo* methods detect footprints across input regions based on their DNase digestion pattern. However, most of these methods were not designed to distinguish between binding sites for specific TFs, and thus cannot automatically label binding sites for specific factors of interest. Motif-centric methods, by contrast, can provide TF-specific prediction, but require pregenerated candidate binding sites for TFs and assess their probability of being TF-bound (active binding sites). Therefore, their performance is limited by the completeness of candidate binding sites provided as they are unable to detect additional regions. Moreover, some of these methods are supervised, meaning they require ChIP-seq data to generate positive and negative training sets, and thus can only be applied to the few TFs with high-quality antibodies. This leads to a major downside as only a minority of TFs have ChIP-seq data available (Wang et al. 2012).

In addition to DNase digestion patterns, more detailed modeling of sequence preference information has been used in TFBSs identification. Hoffman and Birney (2010) have previously proposed a Hidden Markov Model (HMM)-based method, Sunflower, to predict TFBSs based on sequence data alone. Instead of scanning for motif sequences directly, this model takes into consideration the competition between multiple TFs to provide a binding profile for all factors included in the model. While this model maintains the shortcomings of sequence-only methods for identifying TFBSs, it has a greater ability to distinguish which specific TF binds at each predicted TFBS.

Inspired by the success of existing footprinting algorithms and the Sunflower model, we have developed an unsupervised footprinting method, TRACE, based on a HMM framework (Rabiner 1989; Durbin et al. 1998). Our new model integrates both DNase-seq data and PWMs to predict footprints and label binding sites for desired TFs. This method does not depend on pre-generated candidate binding sites or available ChIP-seq data, making it more flexible and broadly-applicable than previous methods.

## Results

### The TRACE Model

TRACE is an HMM-based unsupervised method with the number of hidden states dependent on the numbers and lengths of PWMs included (Fig. 1). The basic structure of our model includes two Background states (the start and end of each open chromatin region), UP, TOP and DOWN states (small peaks surrounding each footprint), and a number of discrete chains of states representing binding sites for each motif included in the model. In addition, the model contains generic footprint states (fp states) representing the regions that have footprint-like digestion pattern but do not match any PWMs in the model (Fig. 1C). Binding site(s) states consist of connected states of its corresponding motif length. For example, the 7-motif CTCF model in Fig. 1C includes state chain of CTCF binding site, 6 additional bait motifs (motif_1, motif_2, …, motif_6) which will be discussed later, and generic footprints whose sequences don’t match any of the motifs included. For each motif included, our model can distinguish its TF-bound from unbound motif sites states, based on their distinct DNase-seq digestion patterns (Supplemental Fig. S1).

**Figure 1.**
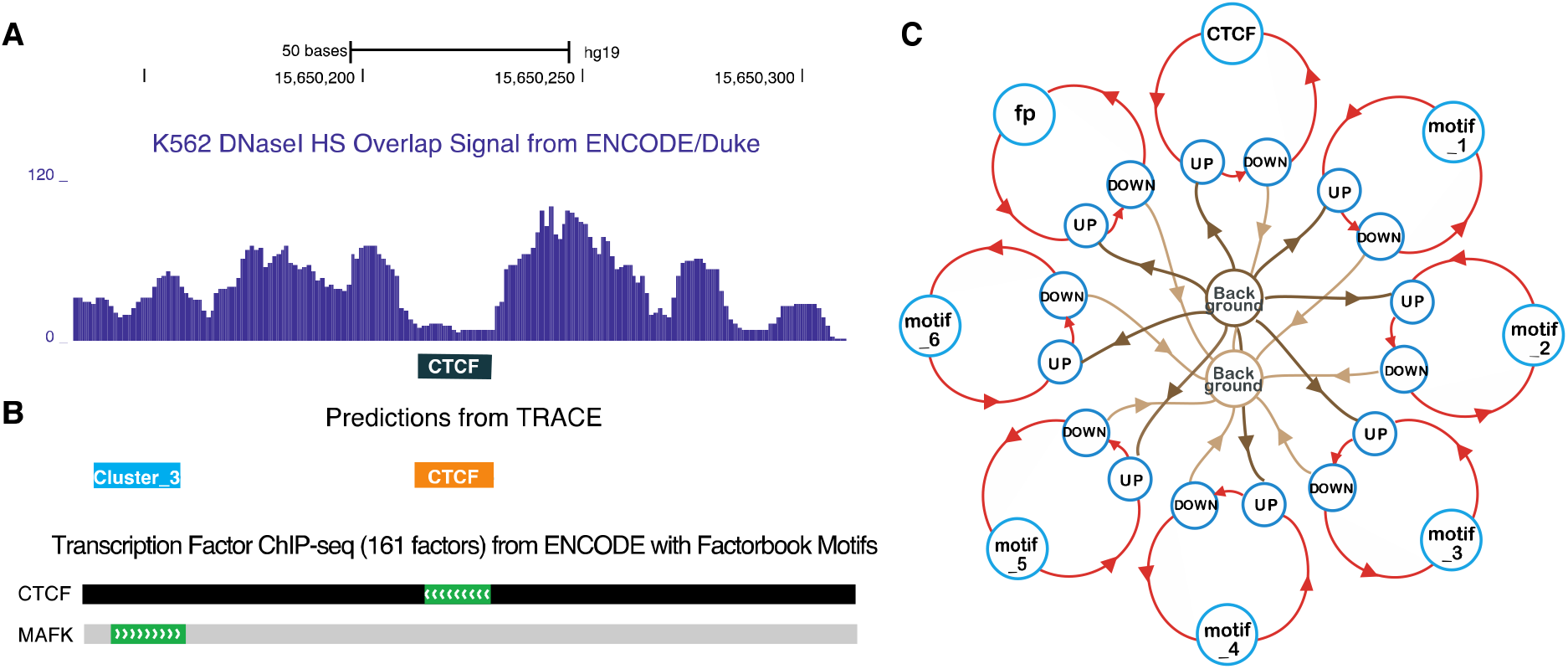
(A) An example of digestion pattern at footprints: DNaseI base overlap signal centered at CTCF motif sites (black box). (B) Predicted binding sites from TRACE using our 10-motif CTCF model match corresponding region of transcription factor binding obtained by ChIP-seq experiments with DNA binding motifs by the ENCODE Factorbook repository. (MAFK is a member of cluster 3 motifs.) (C) Simplified example schematic of a 7-motif CTCF model. Circles represent different hidden states including multiple motifs, lines with arrows represent transitions between different states. For simplicity, TOP states are not shown in the model structure.

TRACE model takes both PWMs and DNase-seq signals as inputs, and models the emission distribution as a multivariate normal distribution using cut count signal and its derivative, and PWM scores at each genomic position. Each binding site (footprint) is expected to be in a region of low sequence density surrounded by a peak of density to either side, with a high PWM score (Fig. 1A, 1B).

### TRACE outperforms existing methods

To evaluate the performance of TRACE relative to published computational footprinting methods, we tested 9 methods (DeFCoM, BinDNase, CENTIPEDE, FLR, DNase2TF, HINT, PIQ, Wellington, as well as a PWM only comparison) on 99 TFs. For a fair comparison across all methods, *de novo* methods were applied to DNase-seq peaks containing the same sets of motif sites that were assessed by motif-centric methods. Receiver operating characteristic curve (ROC) area under the curve (AUC) and Precision-Recall (PR) AUC of predictions of each TF were computed for each method based on the P-values or scores provided, and were ranked across all methods (Fig. 2A, Supplemental Fig. S2, S3).

**Figure 2.**
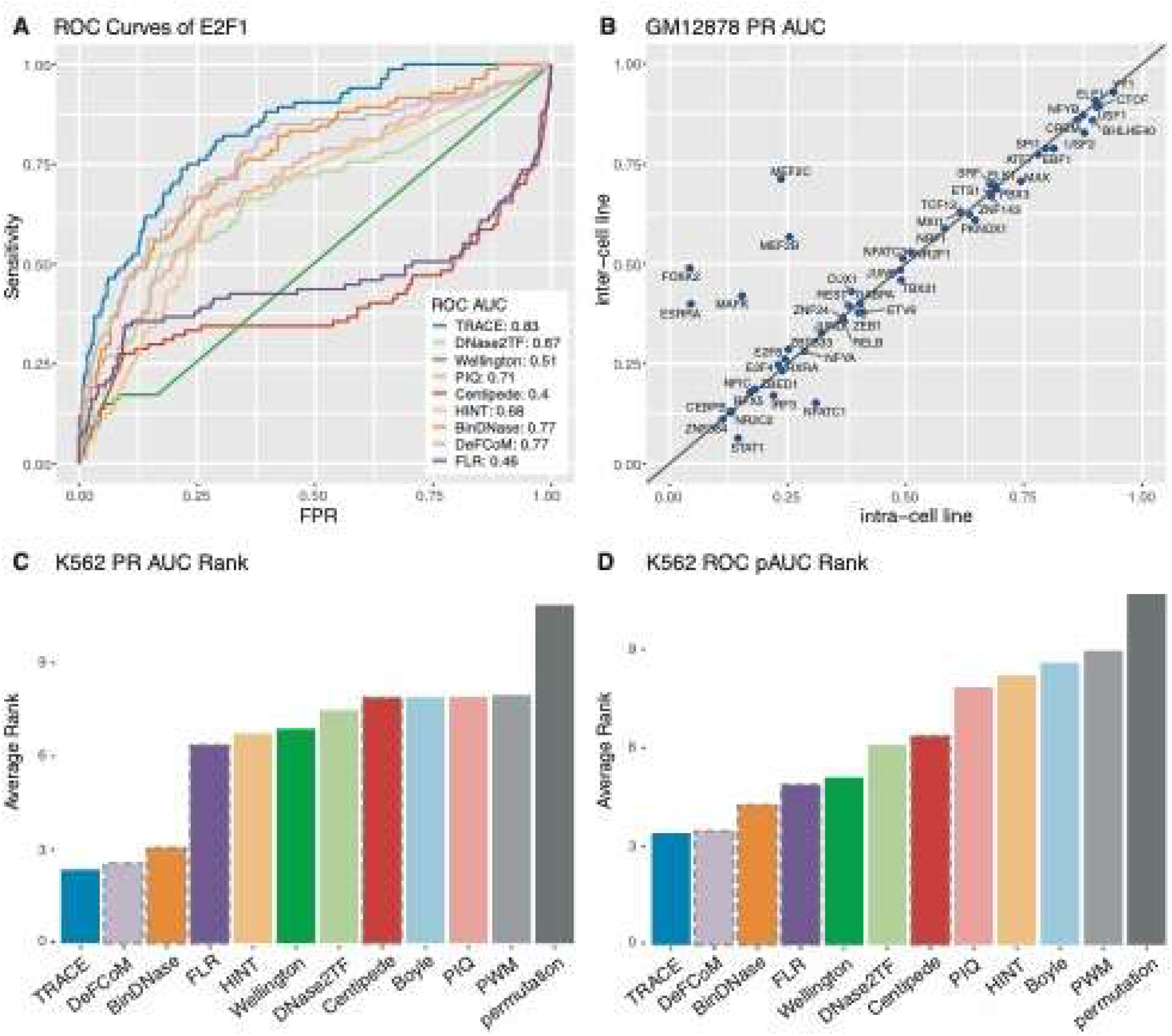
(A) Example ROC curves of E2F1 for all methods evaluated. (B) Cross-cell line comparison of binding sites prediction in GM12878. Each point represents a TF tested, x-axis and y-axis are PR AUCs of applying TRACE using models trained from GM12878 and models trained from K562, respectively. Points above the diagonal line indicate TFs for which inter-cell line model performed better. (C, D) Average rank of PR AUC and ROC pAUC of existing methods across all TFs tested. The Bars with a dashed outline represent motif-centric methods.

Previous studies evaluating computational footprinting methods included a focus on ROC AUC. Although ROC AUC provides a decent measurement for classification performance, the number can be inflated by false positive predictions and can be misleading. For example, the ROC AUC statistic might imply a relatively favorable classification if the method tends to call most samples as positive hits when the data is highly unbalanced, which is the case for many of TFs we tested. Alternatively, we computed partial ROC AUC (ROC pAUC) at a 5% false positive rate (FPR) cutoff. PR AUC was also included in evaluation as it provides a better assessment for false positives. Compared with other footprinting methods, TRACE has the best overall performance based on average rank in both ROC pAUC and PR AUC across 101 tested TFs (Fig. 2). It ranked the first for 25.5% of TFs and in the top 5 for 96.9% of TFs. Compared to other unsupervised methods, TRACE ranked the first for 87.7% of TFs. Notably, TRACE also outperformed supervised approaches including DeFCoM and BinDNase. Therefore, TRACE can predict TF footprints with performance equal or better than the best published methods, while not requiring positive and negative training datasets.

### Bait motifs improve footprinting prediction accuracy

TRACE provides binding site identification for desired TFs at nucleotide resolution. To search for footprint-like regions with a depletion of DNase I cuts surrounded by regions with high numbers of cuts, the basic structure of our model includes hidden states of TFBSs and their surrounding small peaks. By incorporating DNase-seq data and PWMs information, our model is able to detect footprints with an anticipated DNase digestion pattern and matching motifs (Fig. 1B). One important feature of our model is that states for different motifs are independent of each other, enabling its ability to distinctly label binding sites for multiple TFs. In addition, adding extra motifs to the model for a specific TF can potentially increase the accuracy of identifying TF-specific binding sites. The additional motifs introduced in the model work as baits, discouraging prediction of weakly matching sites and introducing competition into the model, thus decreasing the false positive rates (Hoffman and Birney 2010). Importantly, including PWMs with similar sequence preference would not provide useful information and might decrease our model’s ability to distinguish between binding sites of different motifs, To avoid this, we chose only the root motifs from each motif cluster in the JASPAR CORE vertebrates clustering (Khan et al. 2018) and excluded the cluster that contains the TF of interest (Supplemental Methods). In a N-motif model, the root motifs from N-1 clusters with greatest number of occurrences were selected. These N-1 motifs can provide additional information, making the model more sensitive to the TF of interest.

Overall, we found that adding additional motifs to the model yielded significant improvements over our original method, which had a similar HMM structure but did not include motif information (an option now also provided in TRACE) (Boyle et al. 2011). Using a 10-motif model (the TF of interest plus 9 extra motifs), the average PR AUC from TRACE increased by 0.20 (63.1%) over our original method and ROC pAUC improved by 20%.

By comparing models containing different numbers of extra motifs, we found that adding additional TFs can indeed increase the quality of TFBS identification in most cases. However, this was at the expense of considerably increased computational time. Thus we determined that an optimal trade-off between performance increase and computational time was the 10-motif model, which is used in the remainder of this study.

### TRACE can be applied accurately across cell lines

We next performed cross cell-line validation using models trained from K562 DNase-seq data and applied them to GM12878 to test their performance compared to models trained on GM12878. Due to fewer available validation data in GM12878, this comparison was on 52 TFs. Our results indicate that TRACE is able to provide accurate prediction in one cell line using a model trained from another cell line, and that intra-cell line and inter-cell line predictions have comparable overall performance (Fig. 2B). This suggests that our data processing steps successfully capture the signature information of DNase digestion and diminish between-dataset variance to a degree sufficient for effective prediction across cell lines. It also suggests that the DNase digestion pattern of binding sites for most TFs are preserved across cell types.

However, there were some exceptions, for example ESRRA had a much better performance in the inter-cell line test. This TF has far fewer active binding sites in GM12878 (7.6% prevalence) than in K562 (31.3 % prevalence), and so our method may not be able to learn an accurate model from the GM12878 data. This suggests that the model should be trained using the datasets with highest quality and most representative of the true genome-wide binding, after which the same trained model can be applied across all cell types of interest.

TRACE’s cross-cell line application allows fast and large-scale TFBSs prediction using existing models without repetitive model training which is the most time-consuming step. It also shows TRACE’s advantage over supervised methods’ limited usage as only a very small fraction of TFs have ChIP-seq data available (Supplemental Fig. S4). To further showcase this flexibility, we have generated models for 526 JASPAR root motifs and made them available through our github site.

### TRACE can perform as well using ATAC-seq data

ATAC-seq also provides chromatin accessibility information (Buenrostro et al. 2013) and has also been proposed to be used in footprinting analyses. We tested TRACE on ATAC-seq and OMNI-ATAC seq data (Supplemental Methods) to evaluate the performance of our model compared to other models designed to work with this data type. In this evaluation, our results were compared with HINT-ATAC (Li et al. 2019) and DeFCoM, as their original publications included ATAC-seq-based evaluation and showed good performance.

Overall, TRACE maintains the best performance among these three methods as it ranked best for both PR AUC and ROC pAUC (Fig. 3A, Supplemental Fig. S5). We then compared the TRACE prediction accuracy using DNase-seq and ATAC-seq data for each TF in GM12878 (Fig. 3B, Supplemental Fig. S6). This analysis showed that ATAC-seq provides comparable TFBS identification potential as DNase-seq but that TRACE works slightly better with DNase-seq with 60% of TFs showing better PR AUCs when using DNase-seq data. The TFs that showed significant lower PR AUC using DNase-seq were also due to worse

**Figure 3.**
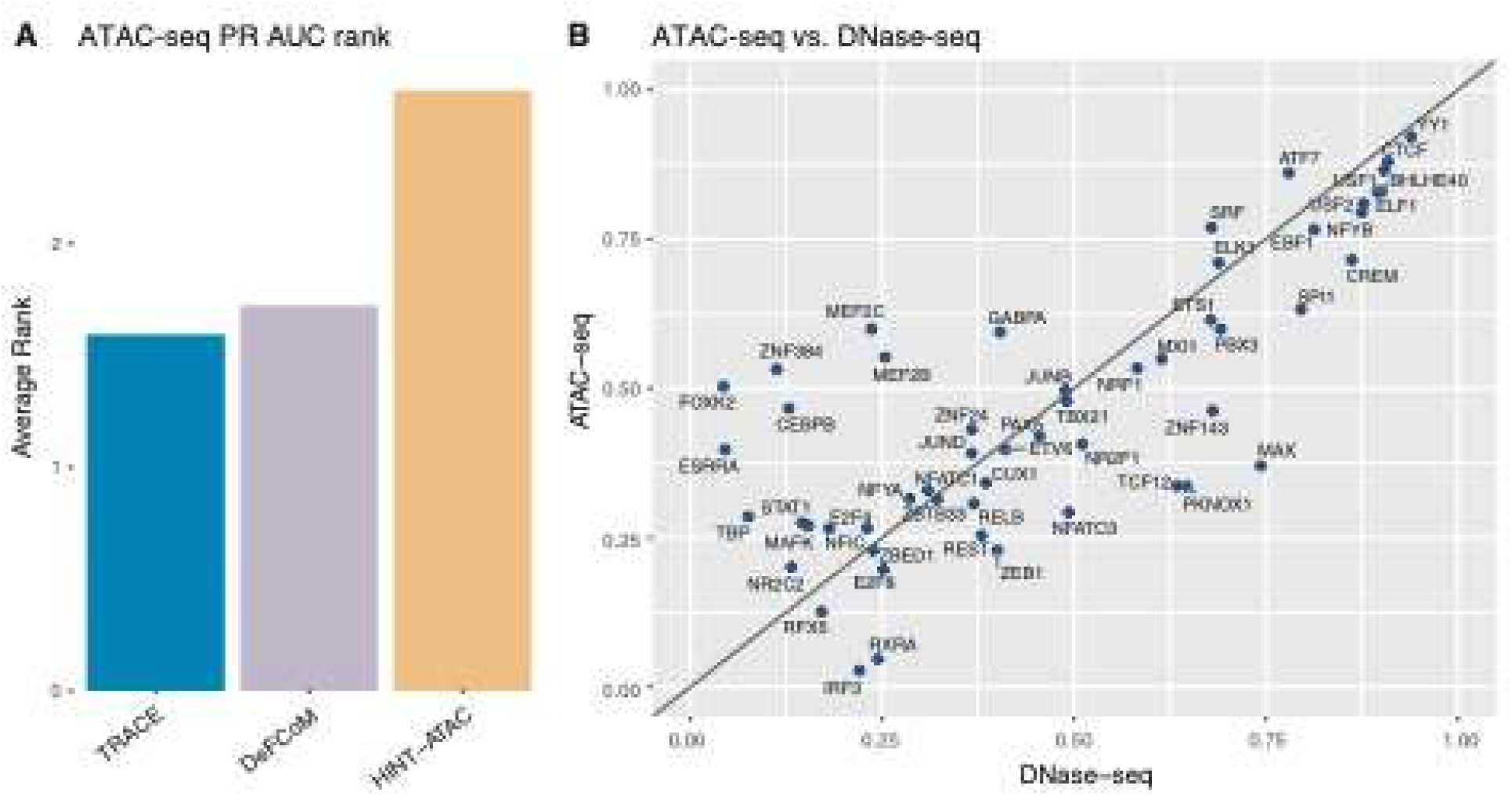
(A) Average rank of PR AUC across all TFs tested using ATAC-seq data for TRACE, DeFCom, and HINT-ATAC. (B) DNase-seq and ATAC-seq based TRACE performance comparison on PR AUC. training data imbalance from GM12878 DNase-seq peak. For example, training sets from ATAC-seq data for FOXK2, ZNF384, CEBPB and TBP all have at least 100% increase of prevalence compared to DNase-seq training sets. To make sure performance difference between these two was not due to the deeper sequencing depth of DNase-seq, we also performed TRACE on a DNase-seq dataset that has comparable and fewer reads than ATAC-seq, and still got similar results (Supplemental Fig. S12). We further downsampled our datasets and found that footprinting performance would drop significantly if read count was below 50 million.

### DNase footprinting has stable performance despite variable levels of data imbalance

It has been noted that not all TFs can have their active binding sites predicted accurately by computational footprinting, regardless of the algorithm applied. Our evaluation of existing footprinting methods indicates that all methods share similar performance trends across all TFs (Fig. 4A left panel). This pattern also exists when assessing candidate binding sites by PWM scores alone (Fig. 4A right panel). In fact, the footprinting performance gain against PWMs is only marginal for some TFs and using PWM scores alone can even outperform all footprinting methods for 2 TFs among the 99 TFs we tested here (Fig. 4B).

**Figure 4.**
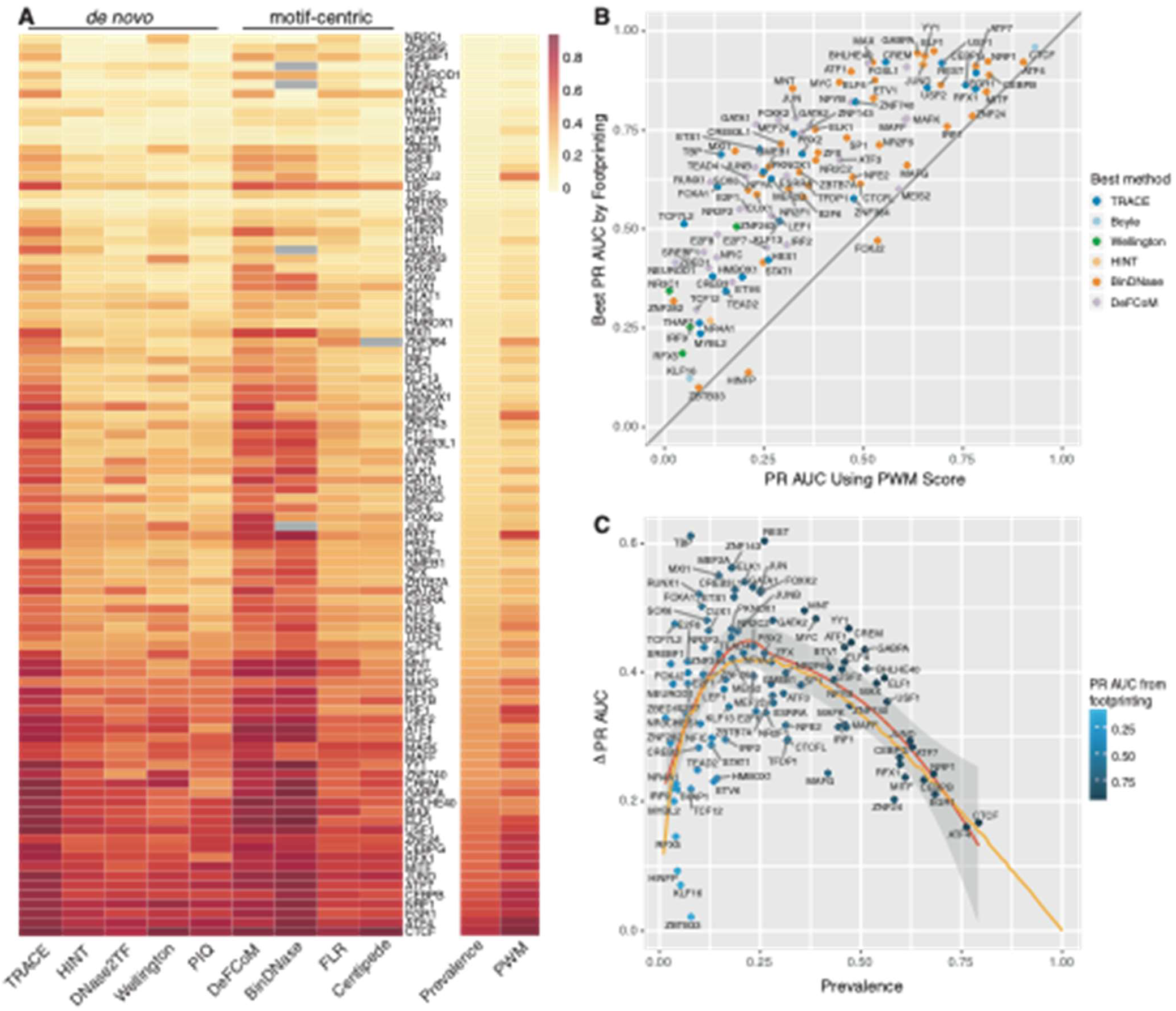
Computational footprinting methods share similar performance patterns. (A) Heatmap of PR AUC of all TFs tested from existing methods sorted by prevalence. (B) Comparison between the best PR AUC among all footprinting methods (y-axis) and PR AUC from using PWM score alone (x-axis) for every TF tested. (C) Performance improvement of footprinting methods over permutation for each TF colored by its best PR AUC from footprinting. Orange line is from a simulation test using positive instances drawn from N(10, 8), and negative instances from N(0, 7) to demonstrate expected PR AUC trend as binding prevalence changes.

We suspect that the poor performance from footprinting might be partially due to the imbalance of positive (P) and negative (N) examples in data sets, as evaluation statistics of prediction for each TF were shown to be associated with its prevalence (fraction of positive samples, P/(P+N), see Methods) (Fig. 4A). Data imbalance affects the quality of model training and, if the data distribution is too skewed, training quality will likely be diminished. We indeed found that some poor performing models were associated with too few positive examples due to their inability to distinguish active and inactive states in model training. However, this only accounts for a small subset and cannot explain the general trend of poor performing TFBSs predictions. Comparing final models for each TF did not reveal significant correlation between prediction accuracy and statistics from different models.

To further explore how computational footprinting may be limited by data imbalance, we compared the best footprinting performance for each TF with a matched-imbalance permutation test of labeled sites (Fig. 4C, Supplemental Fig. S7A, S10). To complement this, simulations were performed with different levels of classification skill and varying imbalance to estimate how PR AUC and ROC AUC values reflect the classifier performance (Supplemental Fig. S7). For the same classification skill, as imbalance changes, we can expect the PR AUC will change correspondingly but ROC AUC and ROC pAUC will stay the same (Saito and Rehmsmeier 2015). However, ROC curves often provide an overly optimistic assessment caused by true negatives used in false positive rate calculation, especially when there is a large skew in the data distribution (Davis and Goadrich 2006).

Instead of comparing AUCs across TFs directly, we measured their performance improvement over random labels (baseline). To examine the general performance gain using computational footprinting, we collected max PR AUCs or ROC AUCs from all existing methods including TRACE and then subtracted the AUCs from the corresponding permutation test. This number was then used as a measurement of footprinting performance advantage over randomly predicted labels. The regression line of PR AUC increase against baseline has a skewed bell-like shape, consistent with the shape of simulated performance generated from a steady model skill (Fig. 4C, Supplemental Fig S7, S10). This suggests that the performance of footprinting is roughly at a stable level and not associated with data imbalance, and a higher evaluation statistic does not necessarily mean a better classification quality for that TF in some cases. Although prevalence may affect evaluation statistic values, we found no evidence that the true classification quality is determined by this data imbalance. Instead, there tends to be a stable level of footprinting classification performance increase compared to random across all TFs.

## Discussion

Incorporating DNase-seq data and PWM information enables TRACE to detect footprints with the desired DNase digestion pattern and matching motifs. By including multiple motifs in the same model, our method provides a better overall TFBS prediction than other existing computational footprinting methods. Since different motifs are treated as separate states in our model, TRACE also has the potential of targeting multiple TFs in a single model. Our method annotates binding sites for the desired TFs across input regions automatically without requiring pre-generated candidate binding sites or additional motif matching steps. In addition, as an unsupervised algorithm, its application is not limited to TFs with available ChIP-seq data.

Although computational footprinting has demonstrated the ability to predict TFBSs at an approximately consistent level, variation in evaluation statistics is still observed across TFs. A previous study showed that not all TFs will leave clear footprint-like nuclease cleavage patterns, and their protection of DNA from cleavage is correlated with residence time (Sung et al. 2014). As a result, for some TFs, footprinting methods might be unable to detect a consistent footprint-like DNase digestion pattern and therefore might fail to label its binding sites correctly. However, there were only limited residence time data available for a small number of TFs and no comprehensive examination on residence time’s impact on footprinting quality has been completed. Although residence time is known to be associated with enzymatic digestion patterns, it is also correlated with the number of active binding sites. GR, AP-1 and CTCF were tested by Sung et al. (2014) as TFs or TF subunits with short, intermediate and long residence time. For those TFs included in our test (NR3C1 as GR group, JUN, JUNB, JUND as AP-1 group, and CTCF), we noticed that TFs with longer residency time tend to have a greater prevalence and a better PR AUC from footprinting (Supplemental Fig. S11). However, neither ROC AUC nor ROC pAUC of these TFs were correlated with residence time. This indicates the possibility that the association between residence time and footprinting ability might be caused by the correlation between performance evaluation statistics and TFBS prevalence. This suggests that the performance disparity may only reflect the changes in fraction of active binding sites among all putative motif sites.

Our evaluation on all footprinting methods indicates that there might be a limited classification accuracy gain that computational footprinting achieves, as the best performance for different TFs all centered at a certain level of classification quality. Our analysis also suggests that evaluation statistics of classification from footprinting may be largely influenced by TFBS prevalence and comparing them directly across TFs may be misleading. Computational footprinting in general might have a maximum potential for how well it can detect TFBSs and only very limited improvement can be achieved beyond this point.

## Methods

### Data and software

DNase-seq data in bam and bed formats and ChIP-seq data in bed format were retrieved from the ENCODE download portal (Supplemental Table S1). The ATAC-seq data for GM12878 cells using standard protocol were obtained from <u>GSE47753</u> (Buenrostro et al. 2013). Omni-ATAC-seq data were obtained from Sequencing Read Archive (SRA) with the BioProject accession PRJNA380283 (Corces et al. 2017). 129 PWMs and cluster information (Supplemental Table S2) were downloaded from JASPAR database (Khan et al. 2018). Motif sites were identified using FIMO (MEME v5.0.3) with default parameters (Grant et al. 2011). Evaluation statistics were generated using python package scikit-learn (Pedregosa FABIANPEDREGOSA et al. 2011).

### Data processing

After bias correction based on model and bias values reported in He et al. (2014), we first counted the number of DNase-seq reads at each location using the 5’ end of the reads, which is the DNase I digestion site. These cut counts were then normalized by non-zero mean of surrounding 10k bp window (within data set normalization) as well as the percentile and standard deviation from the entire region (between data set normalization) (Supplemental Methods). Normalized signals were then smoothed by local regression method R function LOESS (Cleveland et al. 1992) and their derivatives were calculated using savitzky-golay filter using the python package Scipy (Jones et al. 2014). The first derivatives represent the slope of processed signal curve and their signs can indicate the increase or decrease change of data. UP and DOWN states in peak should have positive and negative slopes respectively.

### Evaluation

To assess the performance of TRACE as well as existing computational footprinting tools, we evaluated DeFCoM, BinDNase, CENTIPEDE, FLR, PWM score only, DNase2TF, HINT, PIQ and Wellington based on scores or P-values provided by each method. Candidate binding sites (motif sites) that overlapped with DNase-seq peaks confirmed by ChIP-seq were used as positive set and those not in ChIP-seq peaks but still overlapping DNase-seq peaks made up the negative set. Prevalence was calculated as number of active binding sites (positive set) divided by total number of motif sites (positive set and negative set).

To provide a fair comparison across all methods, we applied *de novo* methods to DNase-seq peaks (with 100bp flanking regions to each side) containing the same sets of motif sites that were included in motif-centric methods tests. For *de novo* methods, only the predictions overlapping with motif sites of tested TFs were included in evaluation; candidate binding sites that were missing from their predictions were also included with an assigned minimum score. For motif-centric methods and PWM only evaluation, only candidate binding sites provided are assessed, so all of their predictions were included in evaluation (Supplemental Methods). Annotations and corresponding scores or P-values were used to calculate the ROC AUC, ROC pAUC at a 5% FPR cutoff and PR AUC values for all TFs.

Permutation test was performed by shuffling labels from footprinting prediction results. Multiple simulation tests were also included based on different levels of positive and negative samples separation and different positive example fractions. Scores for positive and negative groups were randomly drawn from normal distribution of different mean and standard deviation.

## Software availability

TRACE is an open source software; the source code, trained models, and predictions are available on GitHub at https://github.com/Boyle-Lab/TRACE.

## Notes

https://github.com/Boyle-Lab/TRACE

### Competing Interest Statement

The authors have declared no competing interest.

