## Supplemental Methods for "TRACE: transcription factor footprinting using chromatin accessibility data and DNA sequence"

for

### Supplemental Methods

#### 1. Bias-corrections, ~~D~~digestion signals, and derivatives

To generate biased-corrected read counts in a genome with length N $x^B$ (1.1), we first ~~did the~~performed bias correction based on model and bias values reported in He *et al.* (2014). A separate set of raw reads were also used to count raw digestion signal ($x^R$) since they better conserved the overall read counts number difference among different peaks. Only 5' loci were used to count digestion signals in both cases.

$$x^B = (x_1^B, \dots, x_i^B, \dots, x_N^B) \quad (1.1)$$

$$x^R = (x_1^R, \dots, x_i^R, \dots, x_N^R) \quad (1.2)$$

The raw digestion signals were then normalized by the mean of non-zero read counts inside surrounding 10kb window $W$ (1.3) and smoothed using local regression function LOESS from R with degree of the local polynomial = 2 and a small span so that 30 surrounding points were used in the fitting. The resulting signals were saved as read count file that was used in TRACE.

$$x_i^{R \cdot norm} = \frac{x_i^R}{\sum_{j \in W} x_j^R / \sum_{j \in W} 1(x_j^R > 0)} \quad (1.3)$$

We also applied within-dataset normalization (1.4) on the bias corrected signals by surrounding 10kb window, as well as between-dataset normalization (1.5) based on kth percentile $p_L^k$ and standard deviation $\sigma_L$ of the ATAC/DNase-seq peak $L$ or region of interest that contains $x_i$. R function LOESS was then applied on normalized signal $x_i^{B \cdot norm2}$ for local regression.

$$x_i^{B \cdot norm1} = \frac{x_i^B}{\sum_{j \in W} x_j^B / \sum_{j \in W} 1(x_j^B > 0)} \quad (1.4)$$

$$x_i^{B \cdot norm2} = \frac{1}{1 + e^{-(x_i^{B \cdot nor} \quad - p_L^k)/\sigma_L}} \quad (1.5)$$

The normalized and smoothed bias-corrected signals were then used to calculate derivatives using Savitzky-Golay filter from python package Scipy. This method fits data into a second order polynomial with window size of 9. There values were stored in deviation file and used as input data in TRACE.

#### 2. Hidden Markov Model for footprinting

##### 2.1 Model parameters

Hidden Markov models (HMM) have widely applied in bioinformatics. It can represent probability distributions over sequence of observations $O = (O_1, O_2, \ldots, O_T)$ generated by hidden state path $H = (H_1, H_2, \ldots, H_T)$. In TRACE, observations are our processed data and hidden states are different chromatin labels such as footprints and small peaks. Observed features at each genomic position t ($O_t = (O_{t,1}, O_{t.2}, \ldots, O_{t,D})$) include cut count, derivative of ATAC/DNase-seq cut counts and PWM score. HMM parameters also include emission distribution $b_i(k) = P(O_t = k|H_t = i)$, initial states probability $\pi_i = P(H_1 = i)$ and transition probability $a_{ij} = P(H_t = j|H_{t-1} = i)$.

##### 2.2 Model details

Our model was build based on the idea of generalized HMM, in which each motif consists of n states each representing one position in its PWM (n is the length of PWM.). But each state in a motif can only transition to the next state in that motif, and the last state in this motif will transition to state of the small peak, so each motif can still be considered as an individual large state but its parameters at each base pair can be captured separately. There are also fp states representing generalized footprints which do n~~’~~ot match any motif in model. Fig. 1B ~~is just~~ shows this in a simplified structure of TRACE model.

Two Background states represent starting and ending positions for each region of interest. Small peaks that surround footprints are divided into UP, TOP, and DOWN state with one-direction connection. DOWN state will either transit to a footprint state or the end of open chromatin region state (Background state). Only the last state in motif states or start of the region (Background state) can transit to UP state.

To better predict the functional binding sites, we included two sets of states of each motif to represent active and inactive binding sites. For each transcription factor, TRACE will differentiate and predict its functional binding sites and those regions with a matching motif but are no~~n’~~t necessarily bound by that TF.

##### 2.3 Model learning

Given sets of observed features, the Baum–Welch algorithm ~~were~~ was applied to find the maximum likelihood estimate of the parameters $\lambda = (a_{ij}, b_i(k), \pi_i)$. ~~Forward-backward algorithm is applied here.~~ Forward probability $\alpha_t(i)$ (2.1) is the probability of observing the partial sequence $(O_1, O_2, \dots, O_t)$ such that the state $H_t$ is $i$. Backward probability $\beta_t(i)$ (2.2) is the probability of observing the partial sequence $(O_{t+1}, O_{t+2}, \dots, O_T)$ such that the state $H_t$ is $i$.

Forward algorithm induction: (2.1)

- Initialization: $\alpha_1(i) = \pi_i b_i(O_1)$
- Induction: $\alpha_{t+1}(i) = \left[\sum_{j=1}^{N} \alpha_t(j) a_{ji}\right] b_i(O_{t+1})$
- Termination: $P(O|\lambda) = \sum_{j=1}^{N} \alpha_T(j)$

Backward algorithm induction: (2.2)

- Initialization: $\beta_T(i) = 1$

- Induction: $\beta_t(i) = \sum_{j=1}^{N} a_{ij}\, b_j(O_{t+1})\beta_{t+1}(j)$

The Baum–Welch algorithm iteratively updates model parameters $\lambda$. It combines the entire observation $O$ and the forward/backward variables to obtain the probability of being in state $i$ at time $t$ and in state $j$ at time $t+1$ as $\xi_t(i,j)$ (2.4) and the probability of being in state $i$ at time $t$ given the observation sequence as $\gamma_t(i)$ (2.5). $\xi_t(i,j)$ and $\gamma_t(i)$ will then be used for calculation of $\bar{a}_{ij}$ (2.6) and $\bar{b}_i(k)$ (2.7) to update $\lambda$. These newly estimated $\lambda$ will replace the previous $\lambda$ and then to be used in the next iteration, until the increase of $\log P(O|\lambda)$ is smaller than predetermined small number.

$$\xi_t(i,j) = P(H_t = i, H_{t+1} = j|O,\lambda) = \frac{\alpha_t(i)a_{ij}b_j(O_{t+1})\beta_{t+1}(j)}{P(O|\lambda)} = \frac{\alpha_t(i)a_{ij}b_j(O_{t+1})\beta_{t+1}(j)}{\sum_{i=1}^{N}\sum_{j=1}^{N}\alpha_t(i)a_{ij}b_j(O_{t+1})\beta_{t+1}(j)} \quad (2.3)$$

$$\gamma_t(i) = P(H_t = i|O,\lambda) = \sum_{j=1}^{N}\xi(i,j) \quad (2.4)$$

$$\bar{a}_{ij} = \frac{\sum_t \xi_t(i,j)}{\sum_t \gamma_t(i)} \quad (2.5)$$

$$\bar{b}_i(k) = \frac{\sum_{t,O_t=k}\gamma_t(i)}{\sum_t \gamma_t(i)} \quad (2.6)$$

The emission distribution ($b_i(k)$) (2.1) for each hidden state $H_t = i$ is modeled with a ~~multivariance~~ multivariate normal distribution, with D-dimensional mean vector $\mu_i = (\mu_{i,1}, \mu_{i,2}, \dots, \mu_{i,D})$ and full covariance matrix $\Sigma_i$. D is number of features included as input data. Emission parameters that are estimated in Baum-Welch algorithm include $(\mu_i, \Sigma_i)$ for each state $i$.

$$b_i(k) = P(O_t = k|H_t = i) = P(k|\mu_i,\Sigma_i) = \frac{1}{\sqrt{(2\pi)^D|\Sigma_i|}} e^{-\frac{1}{2}(k-\mu_i)^T(\Sigma_i)^{-1}(k-\mu_i)} \quad (2.7)$$

$$\mu_i = \frac{\sum_{t=1}^{T} O_t \mathbf{1}(H_t=i)}{\sum_{t=1}^{T}\mathbf{1}(H_t=i)} \quad (2.8)$$

$$\Sigma_i = \frac{\sum_{t=1}^{T}(O_t-\mu_i)^T(O_t-\mu_i)\mathbf{1}(H_t=i)}{\sum_{t=1}^{T}\mathbf{1}(H_t=i)-1} \quad (2.9)$$

The Viterbi algorithm was~~ere~~ implemented to find the best hidden path $H$ that maximizes the likelihood $P(H|O,\lambda)$ from observation $O$ and estimated model $\lambda$. It finds the most probable hidden path $H$ recursively by ~~getting~~ determining $\delta_t(i) = \max_{1 \le i \le N} P(H, O|\lambda)$, the highest probability path ending in state $i$, then backtrack to obtain the best path. Since one of the features in TRACE model is PWM score, an optional bit score threshold can be included to reduce the false positive calls. When threshold is provided, $\delta_t(i)$ for state i at position t will be set to a minimal value if PWM score at t is below the threshold.

**3. Bait motif selection**

~~Besides~~ In addition to the states of the TF of interest, the TRACE model also includes other motifs which serve as bait motifs. Adding bait motifs can potentially reduce the false positive labels of regions with footprint-like digestion patterns but a weak sequence match with the TF of interest, as those regions might have higher sequence preference for bait motifs. This binding competition can potentially increase the accuracy of identifying binding sites.

In order to add useful information in the model, the extra motifs should not have similar binding preference, otherwise they will only contain repetitive sequence information and be treated as same states by TRACE. To ensure all motifs included in the model are different, we obtained hierarchical clustering information of PFMs from JASPAR database using the RSAT matrix-clustering tool (Castro-Mondragon et al. 2017). The motifs at the root of each tree encompass~~es~~ all the position-specific scoring matrices (PSSMs) of a cluster. Those root alignments are the only PWMs should be added in TRACE model as baits. The root motif from the cluster that contains the TF of interest should also be excluded from the model. To decide which motifs should be added in the model, we scanned each root motif across the genome and ranked their numbers of occurrences. For a N-motif model for a certain TF, the bait motifs will be (N-1) most abundant root motifs from the clusters that don't contain that TF of interest.

#### 4. Existing footprinting methods evaluation

To assess the performances of existing computational footprinting tools including DeFCoM, BinDNase, CENTIPEDE, PWM score only, DNase2TF, HINT, FLR, CENTIPEDE, PIQ, Wellington and original Boyle method, we followed the motif-centric evaluation approach and tested on chr1. For motif-centric methods, candidate binding sites overlapped with DNase-seq peaks were used as testing sites and included in evaluation. For *de novo* methods, that same group of DNase-seq peaks were selected as testing regions.

For evaluation, all sites and their scores from motif-centric methods are included and assessed. For *de novo* methods, only those predicted footprints overlapped with at least 10% of a motif site were included; missing candidate binding sites from those input DNase-seq peaks were also included and assigned with a minimum score. We used the default settings for the majority of existing methods. Those ones with modifications or needing clarifications are listed below:.

DeFCoM and BinDNase

Since they are supervised methods, we trained model on chr19 and tested on chr1. Motif sites overlapped with DNase-seq peaks in chr1 and chr19 were used as testing set and training set respectively. We used default parameters as specified by Kähärä and Lähdesmäki (2015) and Quach and Furey (2016).

CENTIPEDE

We utilized cut signal, PWM score, distance to closest TSS and conservation score as input data. The other parameters were set the same as described by Pique-Regi *et al.* (2011).

HINT

We applied HINT on only DNase-seq or ATAC-seq data with default setting, and included the bias correction step. Also, we used newest version of HINT-ATAC for tests involving ATAC-seq data.

Wellington

We used data from double-hit DNase-seq protocol as required by Wellington and set -fdrlimit to 0. For evaluation, the absolute value of its reported scores for each footprint were used.

Boyle method

We have a build-in option in TRACE program to apply Boyle method. We included read counts and slope data and build a 6-state model similar to the model described in Boyle *et al.* (2011), then used the posterior probabilities in evaluation.

PWM score only

We calculated PWM scores for all motif sites that were included in motif-centric methods evolution. This includes only PWMs that overlap open chromatin sites in the current data set. Only PWM scores were used in this evolution.

**5. ATAC-seq pipeline**

The ATAC-seq data for GM12878 were obtained from GSE47753, Omni-ATAC-seq data were obtained from Sequencing Read Archive (SRA) with the BioProject accession PRJNA380283.

These data were processed following Kundaje lab's ATAC-seq pipeline.

(https://github.com/kundajelab/atac_dnase_pipelines)

#### Supplemental Figures

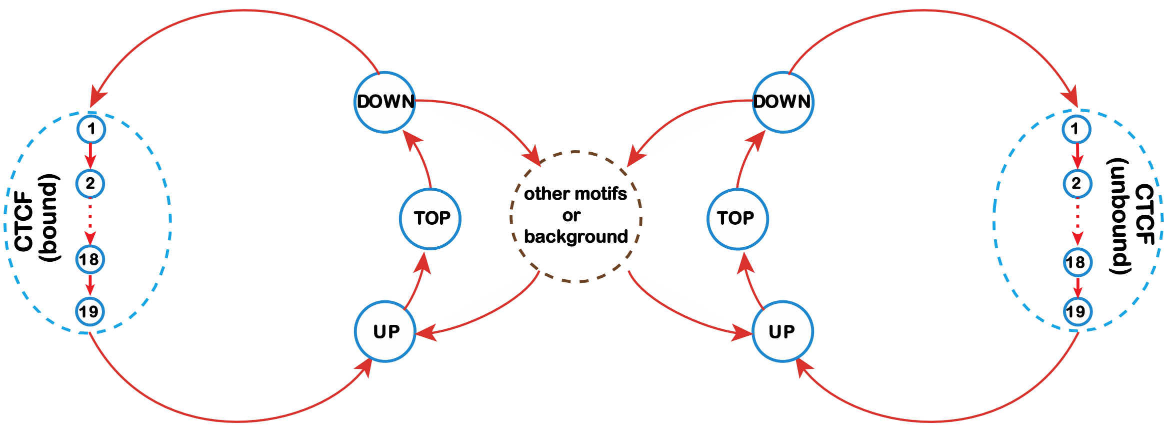


Supplemental Figure 1. Detailed schematic of bound and unbound CTCF state in CTCF model. Circles represent different hidden states including binding sites and peaks, lines with arrows represent transitions between different states. For simplicity, all other motifs, generic footprint states and background are represented by a dashed line circle.

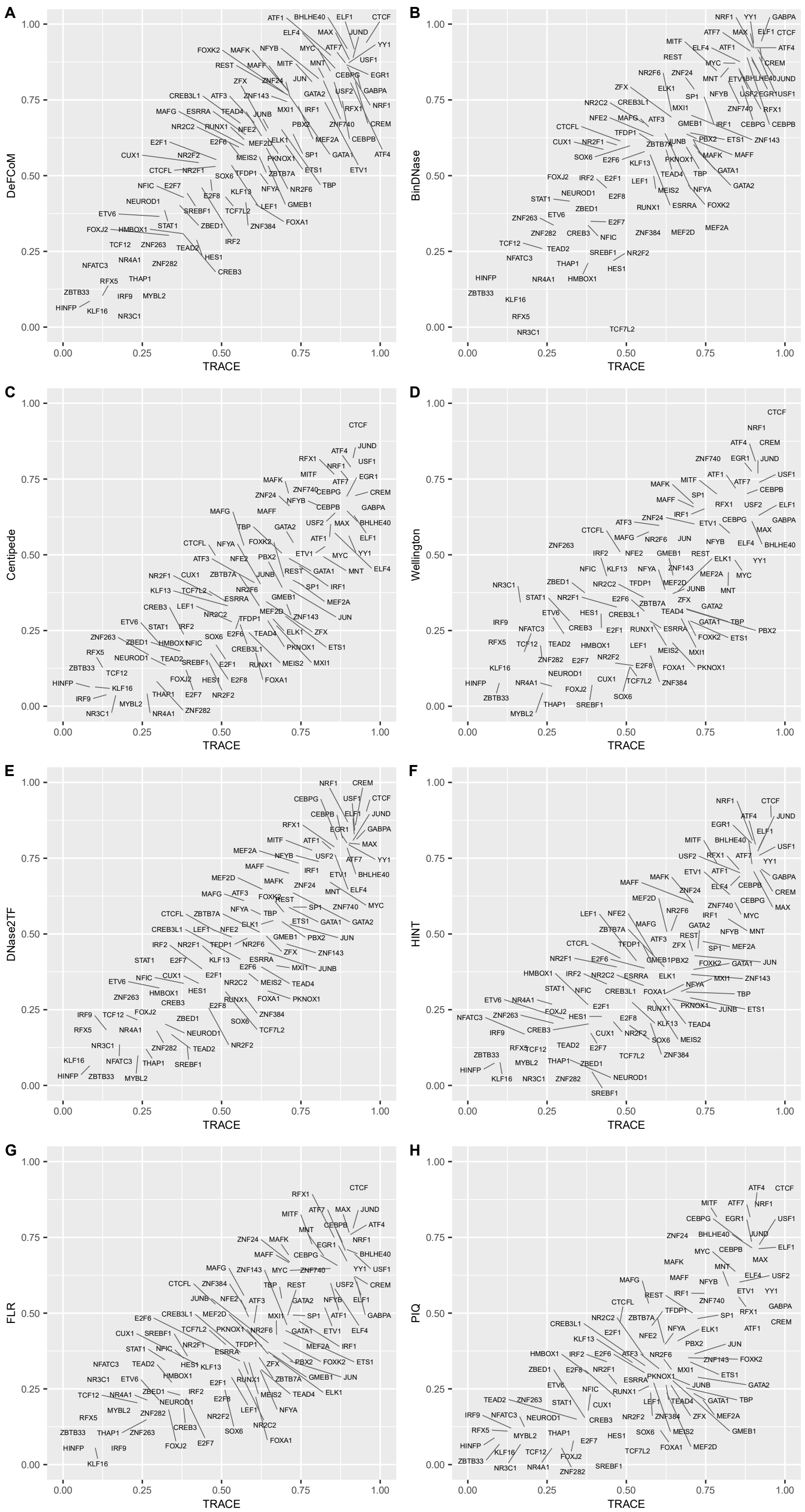


Supplemental Figure 2. PR AUC comparison between TRACE and existing methods for binding sites prediction of all TFs tested

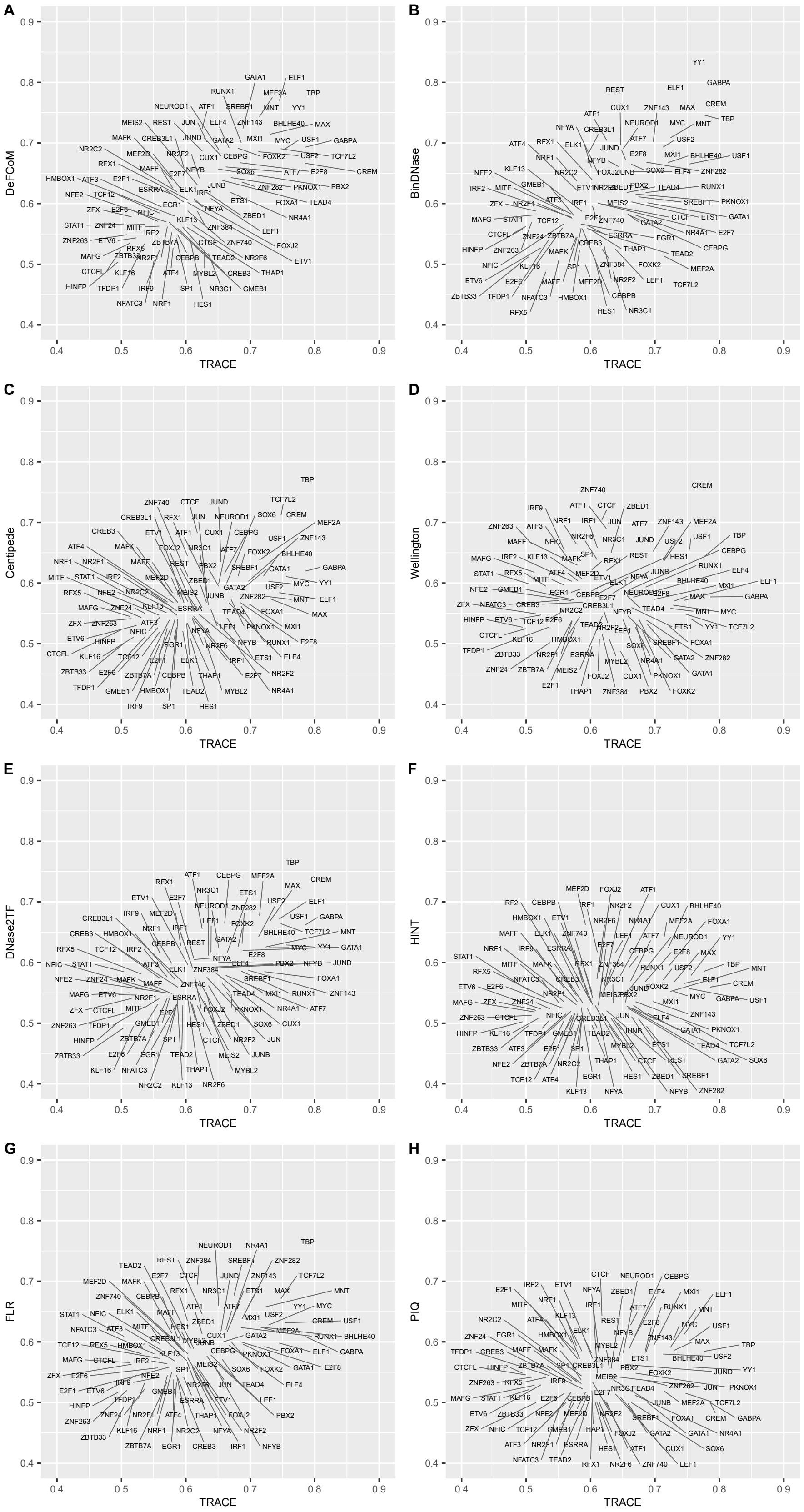


Supplemental Figure 3. ROC pAUC comparison between TRACE and existing methods for binding sites prediction of all TFs tested

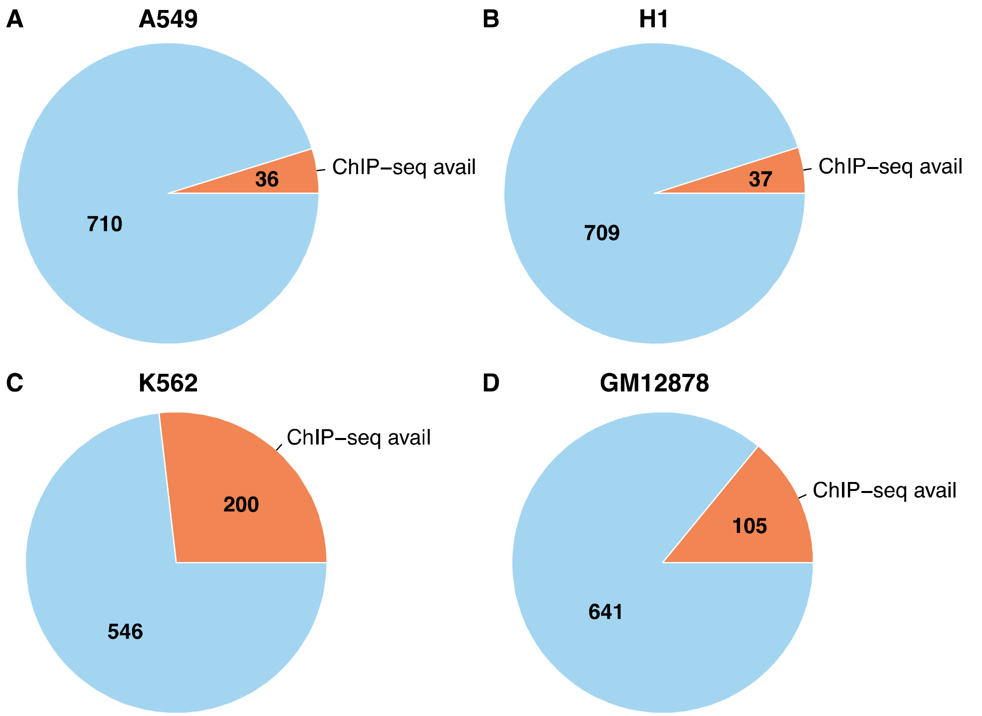


Supplemental Figure 4. Numbers of TFs with ChIP-seq data available in (A) A549, (B) H1, (C) K562 and (D) GM12878 among all motifs in JASPAR CORE database (non-redundant). Each pie plot has a total number of 746 TFs, orange protion represents number of TFs has ChIP-seq data in that cell line in ENCODE.

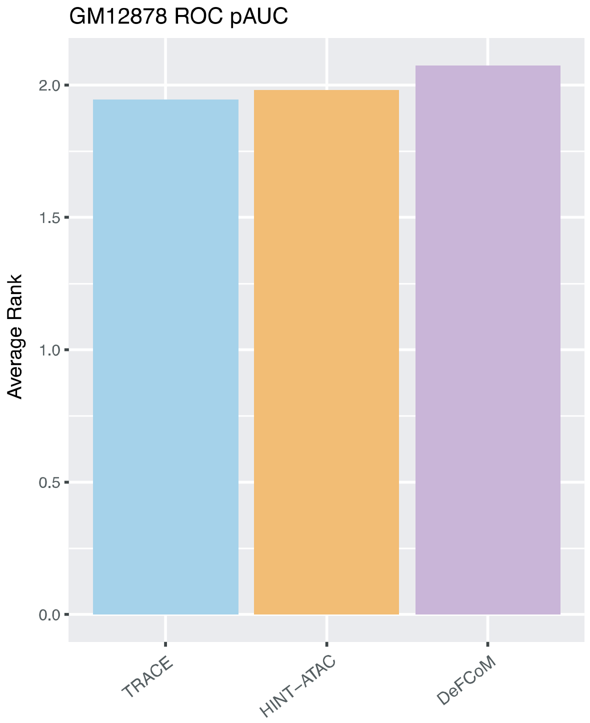


Supplemental Figure 5. Average rank of ROC pAUC across all TFs tested using ATAC-seq data for TRACE, DeFCoM and HINT-ATAC

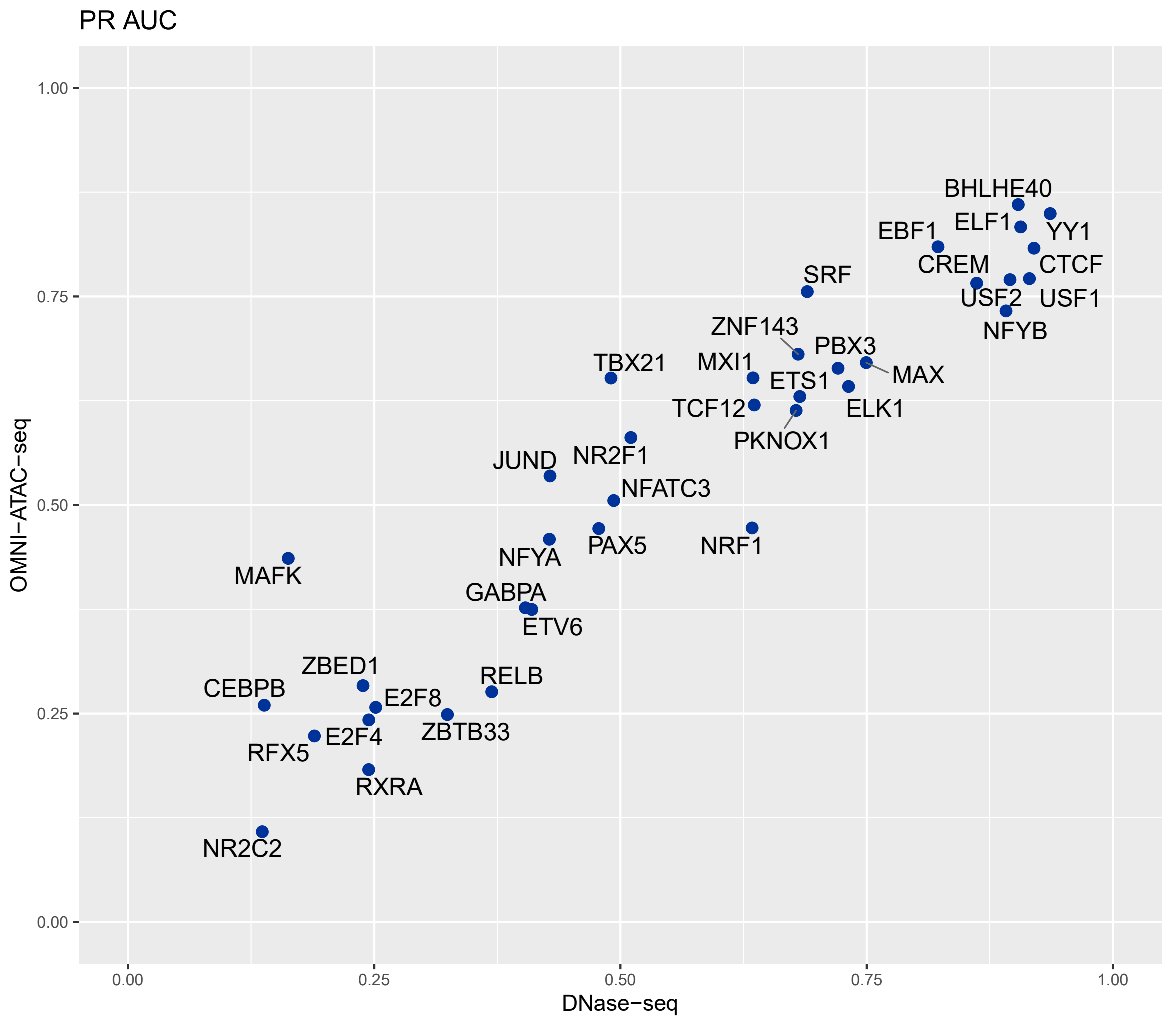


Supplemental Figure 6. DNase-seq and OMNI-ATAC-seq based TRACE performance comparison on PR AUC.

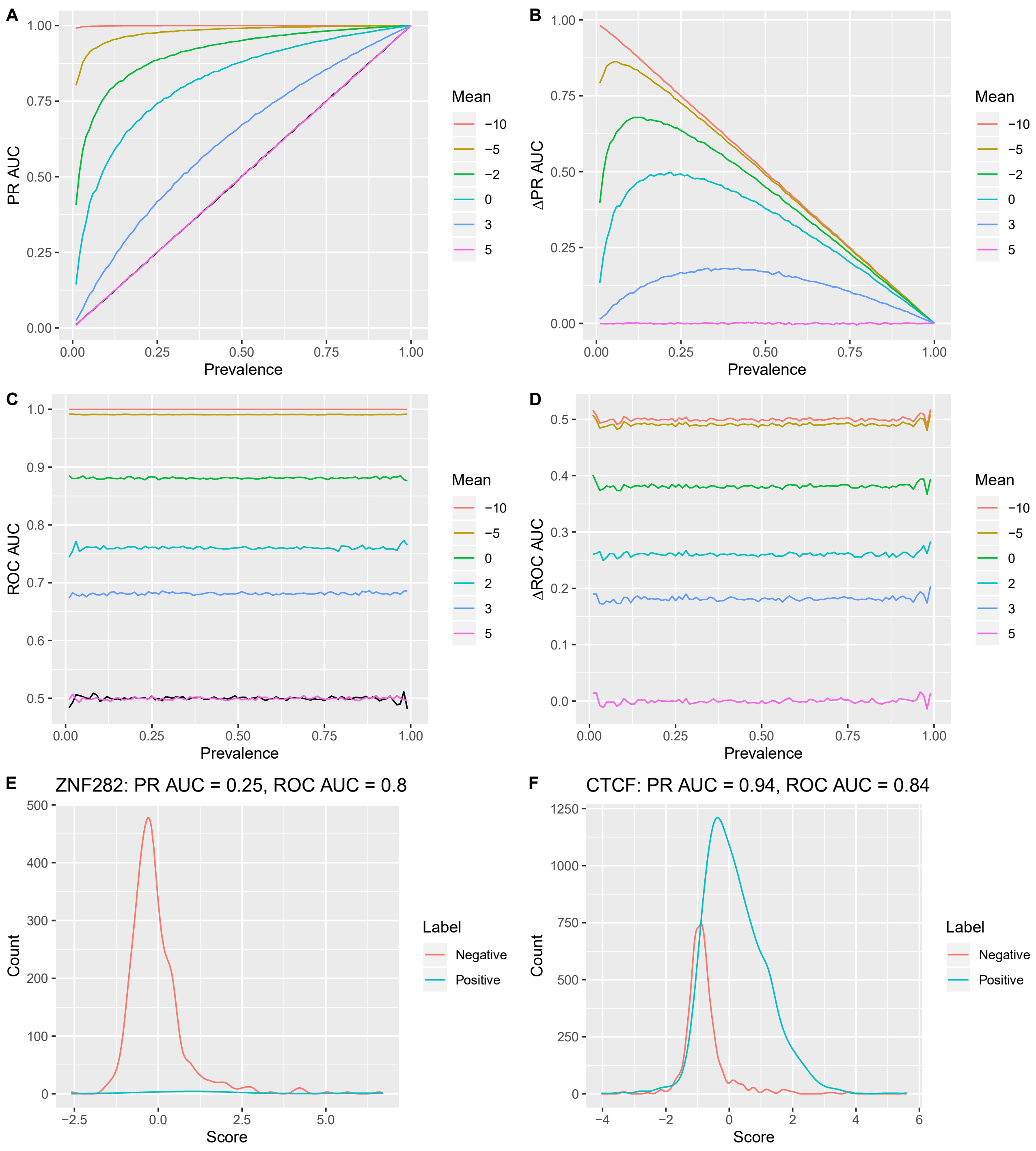


Supplemental Figure 7. (A) PR AUC and (C) ROC AUC from simulation test with different prevalence. Scores for positive examples for all simulation were drawn from the same distribution N(5, 3). Each line represents a different negative example distribution with their own mean value, varying from 5 to -10. Black lines represent AUCs from random labels. (B) PR AUC increase and (D) ROC AUC increase from simulation test with different prevalence. (E, F) Score distributions for ZNF282 and NR2C2 as examples of TFs with different level of data imbalance.

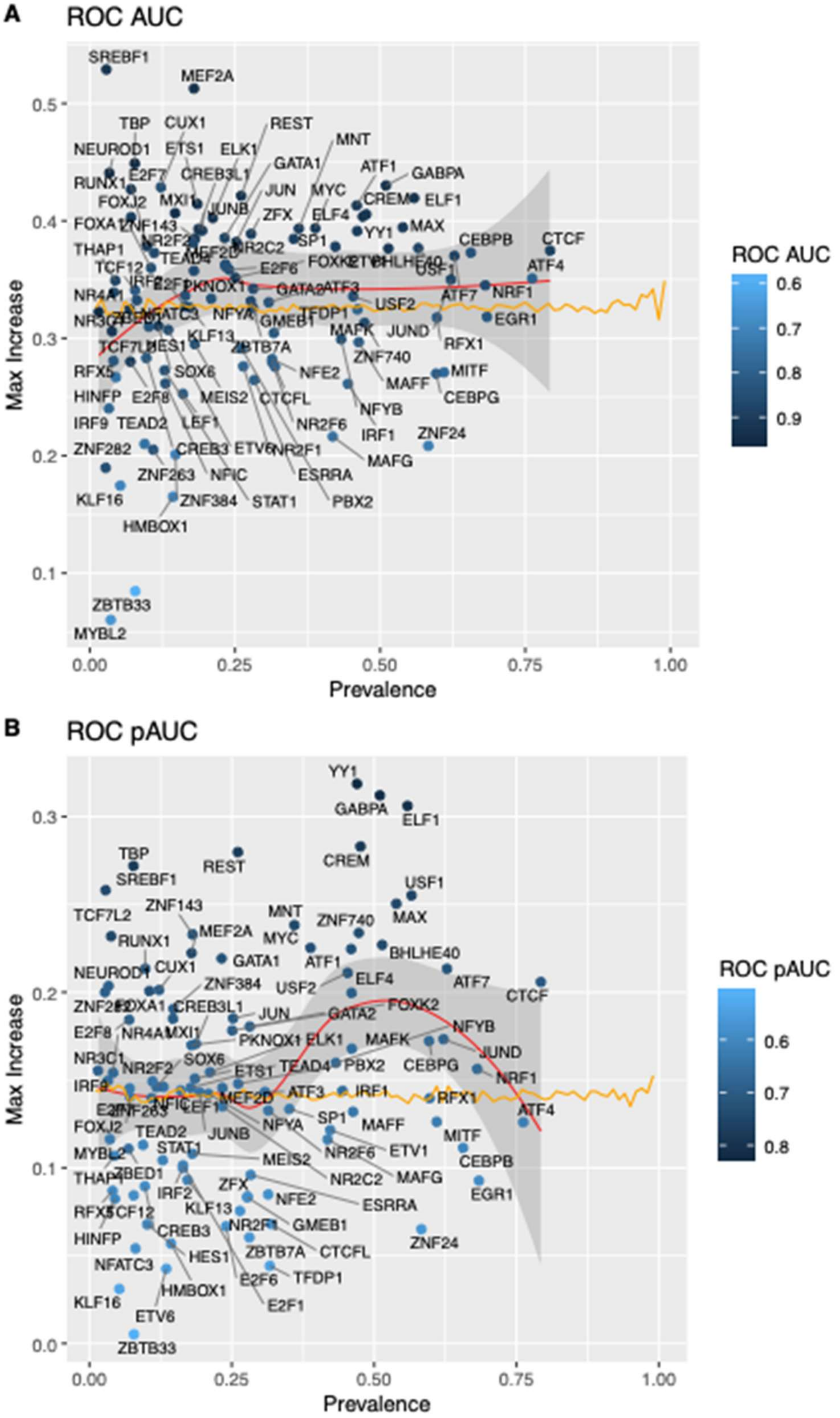


Supplemental Figure 8. increase of footprinting methods' best (A) ROC AUC and (B) ROC pAUC over permutation are not correlated with prevalence. Orange line is from simulation test using positive set from N(10, 8), negative set from N(0, 7).

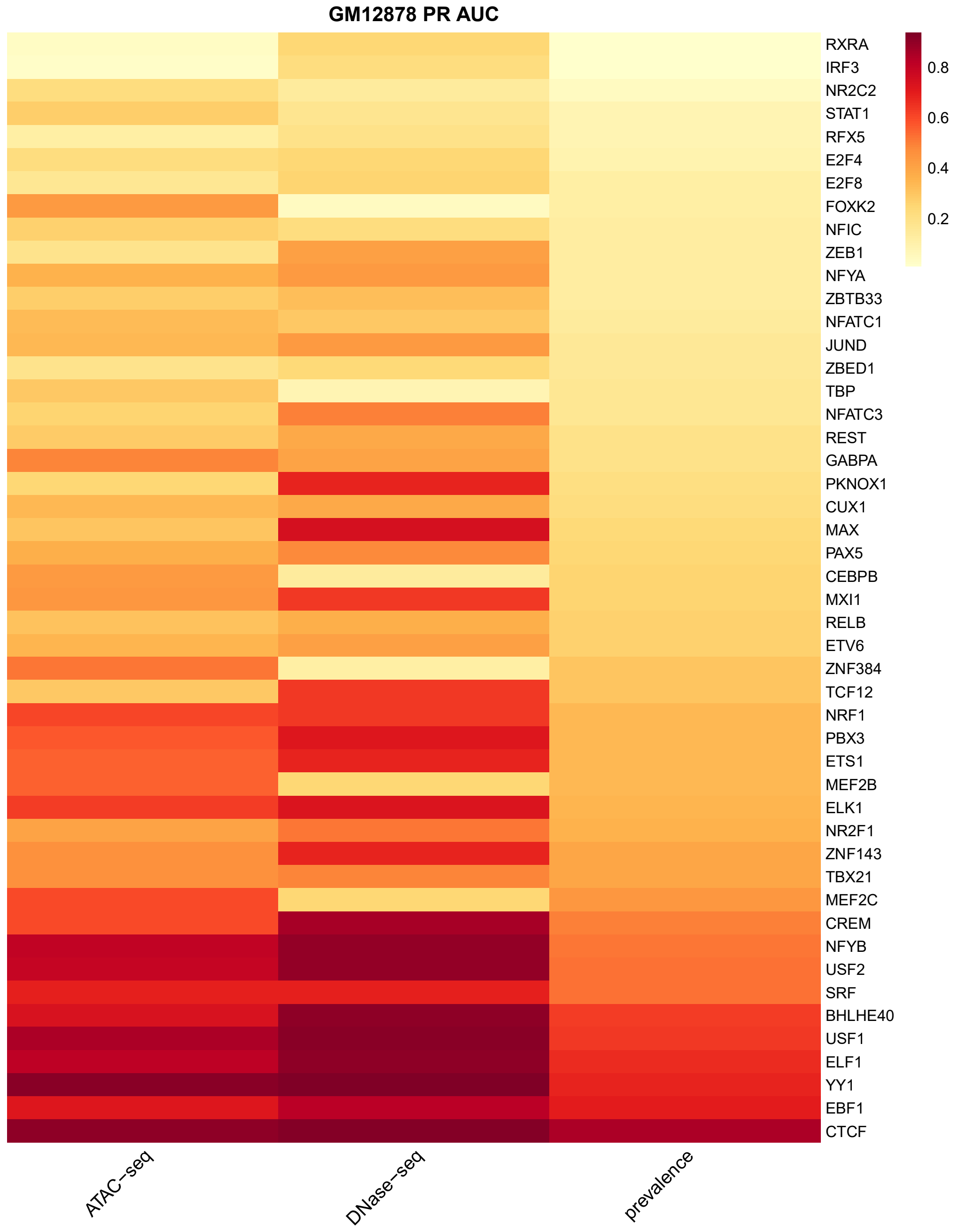


Supplemental Figure 9. Heatmap of PR AUC of all TFs tested using DNase-seq and ATAC-seq data, sorted by prevalence.

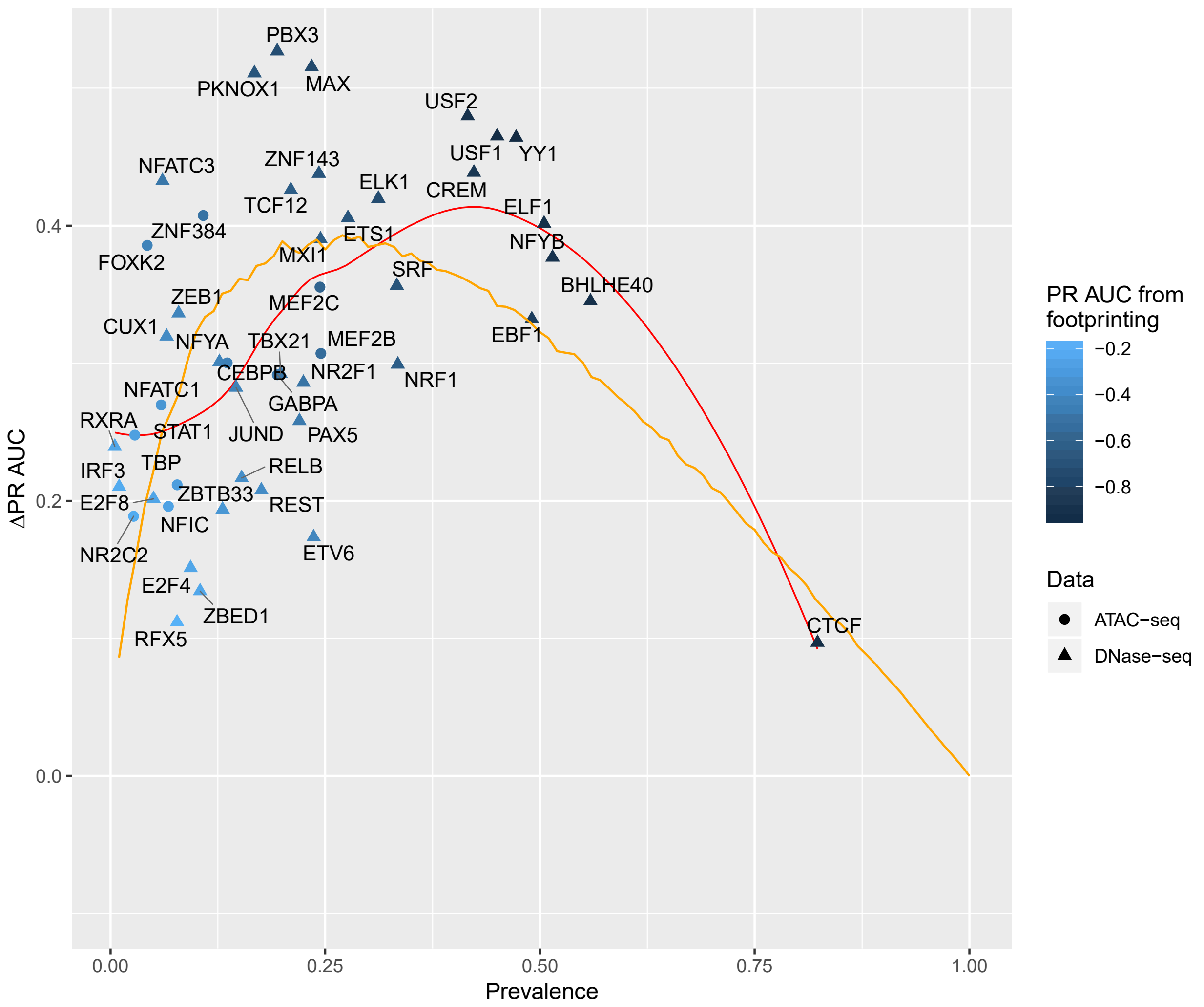


Supplemental Figure 10. Performance improvement of TRACE model over permutation for each TF in GM12878, colored by its best PR AUC from DNase-seq or ATAC-seq data. Orange line is from simulation test using positive instances drawn from N(12, 9), and negative instances from N(0, 9) to demonstrate expected PR AUC trend as binding prevalence changes.

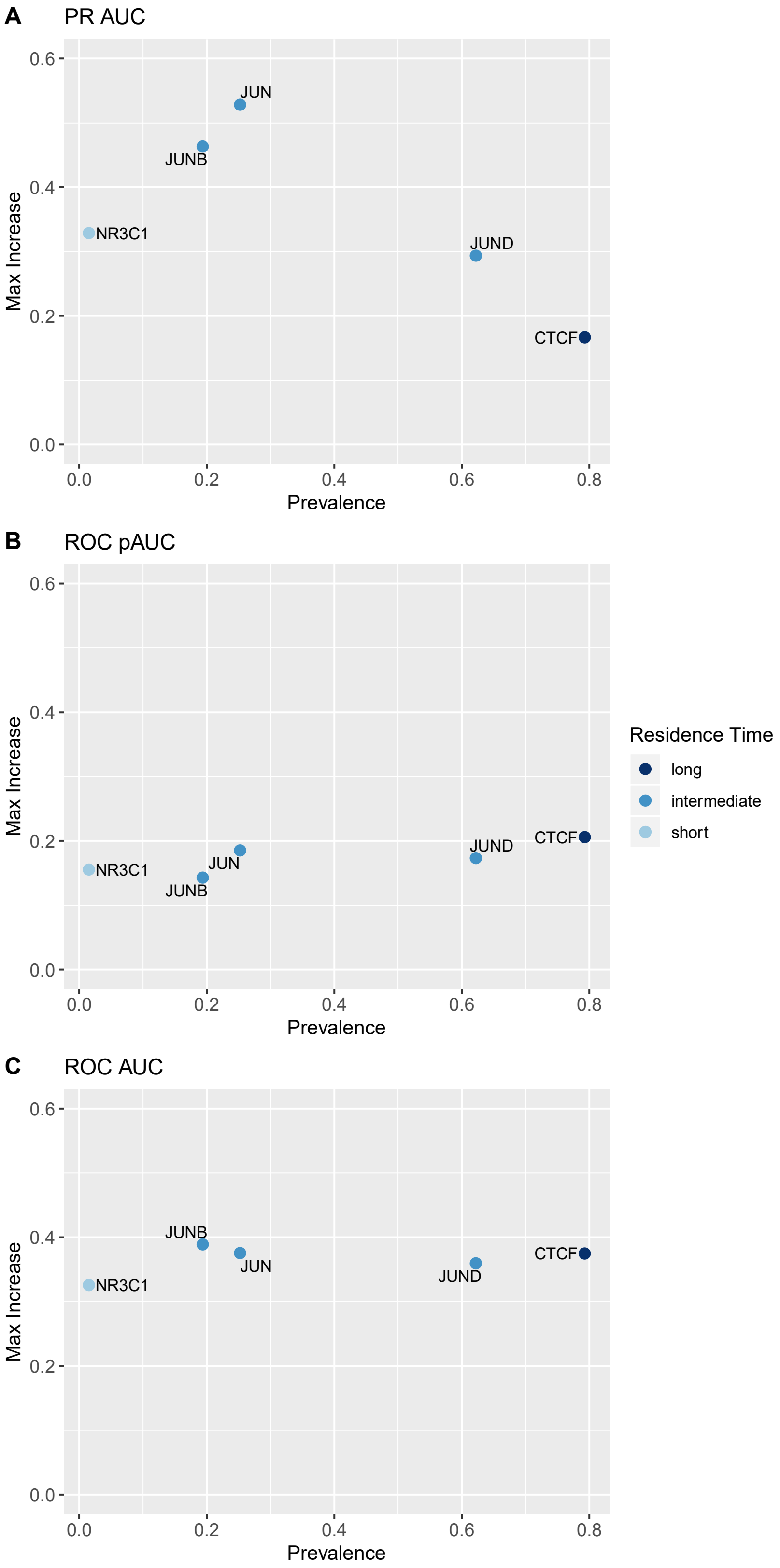


Supplemental Figure 11. Comparison of footprinting performance increase on TFs with (A) short, (B) intermediate and (C) long residence time.

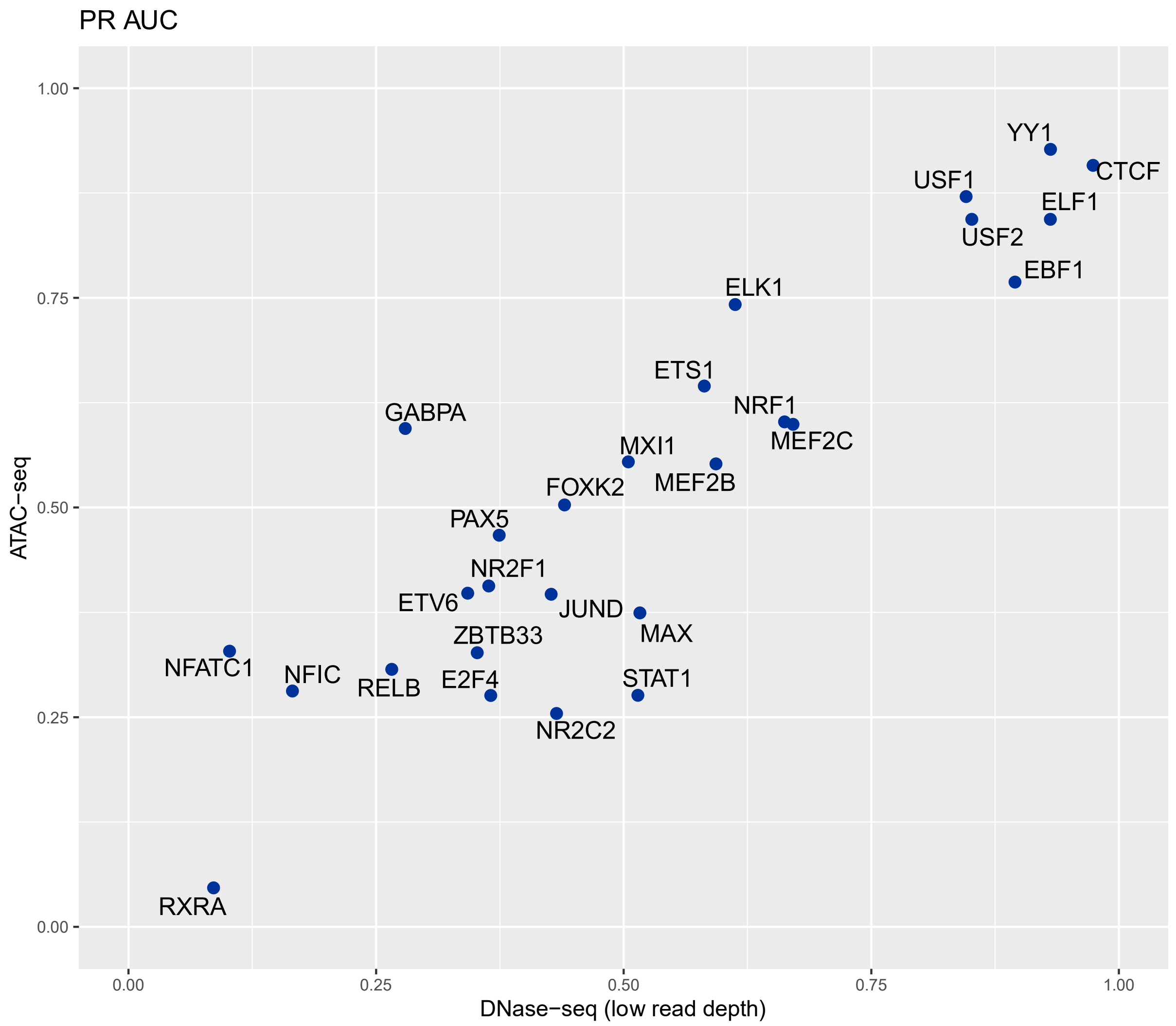


Supplemental Figure 12. TRACE performance comparison on DNase-seq and ATAC-seq with comparable level of read depth. DNase-seq data used in this analysis has fewer reads than the one in Figure 3.

#### Supplemental Table

(A)

| | 1-motif model | | | 10-motif model | | | |
|---|---|---|---|---|---|---|---|
| Size of training set (kilobases) | Number of states | Computational time | Memory | Number of states | Computational time | Memory | Number of cores |
| 71.9 | 28 | <1min | 0.21G | 310 | 4min | 1.4G | 40 |
| 180.6 | 34 | 1min | 0.59G | 316 | 9min | 3.5G | 40 |
| 238.3 | 52 | 2min | 0.98G | 334 | 17min | 4.8G | 40 |
| 883.1 | 32 | 4min | 2.8G | 296 | 62min | 15.6G | 40 |
| 1308.6 | 48 | 7min | 4.2G | 316 | 90min | 20.2G | 40 |

(B)

| | 1-motif model | | | 10-motif model | | | |
|---|---|---|---|---|---|---|---|
| Size of testing set (kilobases) | Number of states | Computational time | Memory | Number of states | Computational time | Memory | Number of cores |
| 71.9 | 28 | <1s | 0.2G | 310 | 3s | 1.1G | 40 |
| 180.6 | 34 | <1s | 0.5G | 316 | 7s | 2.9G | 40 |
| 238.3 | 52 | 1s | 0.9G | 334 | 9s | 4.0G | 40 |
| 883.1 | 32 | 2s | 2.3G | 296 | 29s | 12.8G | 40 |
| 1308.6 | 48 | 4s | 3.5G | 316 | 39s | 16.8G | 40 |

Supplemental Table 3. Computational time and memory TRACE requires for (A) training and (B)Viterbi step, with different sizes of model and training set. Each row shows the time and memory needed to finish a TRACE run for an example TF on a certain number of kilobase length of training data, using models with different numbers of hidden sates depending on motif size. CPU: Intel Xeon E5-2696 v4 @ 3.7GHz
